# BIN overlap confirms transcontinental distribution of pest aphids (Hemiptera: Aphididae)

**DOI:** 10.1101/705889

**Authors:** Muhammad Tayyib Naseem, Muhammad Ashfaq, Arif Muhammad Khan, Akhtar Rasool, Muhammad Asif, Paul D.N. Hebert

**Author notes:** Corresponding author: Muhammad Ashfaq, Centre for Biodiversity Genomics & Department of Integrative Biology, University of Guelph, Guelph, ON, N1G 2W1, Canada.

## Abstract

DNA barcoding is highly effective for identifying specimens once a reference sequence library is available for the species assemblage targeted for analysis. Despite the great need for an improved capacity to identify the insect pests of crops, the use of DNA barcoding is constrained by the lack of a well-parameterized reference library. The current study begins to address this limitation by developing a DNA barcode reference library for the pest aphids of Pakistan. It also examines the affinities of these taxa with conspecific taxa from other geographic regions based on both conventional taxonomy and Barcode Index Numbers (BINs). A total of 809 aphids were collected from 123 plant species at 87 sites across Pakistan. Morphological study and DNA barcoding allowed 774 specimens to be identified to one of 42 species while the others were placed to a genus or subfamily. The 801 sequences obtained from these specimens were assigned to 52 BINs whose monophyly were supported by neighbor-joining (NJ) clustering and Bayesian inference. The 42 species were assigned to 41 BINs with 38 showing BIN concordance; one species (*Rhopalosiphum padi*) was assigned to two BINs, while two others (*Aphis affinis, Aphis gossypii*) were assigned to the same BIN, while one species (*Aphis astragalina*) lacked a qualifying sequence. The 42 Linnaean species were represented on BOLD by 7,870 records from 69 countries. Combining these records with those from Pakistan produced to 60 BINs with 12 species showing a BIN split and three a BIN merger. Geo-distance correlations showed that intraspecific divergence values for 18 of 37 species were not affected by the distance between populations. Forty four of the 52 BINs from Pakistan had counterparts in 73 countries across six continents, documenting the broad distributions of pest aphids.

## Introduction

Although aphids (Hemiptera: Aphididae) are important plant pests, their life stage diversity and phenotypic plasticity have constrained the development of effective taxonomic keys [1,2]. With over 4,700 described species, the Aphididae is the largest family within the Aphidoidea [3]. Most pest aphids belong to the subtribe Aphidina which includes 670 described species [3,4]. Nearly 100 aphid species have been listed as serious agricultural pests; they attack more than 300 plant species [5,6], and lower crop yield by direct feeding and by transmitting viral diseases [7].

Sibling species complexes are common in many pest aphids [8]. Very often, these species are morphologically identical but genetically distinct [9]. They often include anholocyclic biotypes (=clones) with differing host preferences and varying competency for disease transmission [10,11]. Species identification is so challenging that taxonomic keys are either ineffective or only useful for a particular geographic area or taxonomic group 12]. These deficits have prompted the search for alternative approaches for identification such as cybertaxonomy [13] and DNA sequencing [14,15]. However, the later approach has gained stronger uptake due to its universal applicability, low cost, and strong performance [16].

Past studies have shown that DNA-based approaches can enable both specimen identification and the clarification of cryptic species complexes [17,18]. Diverse mitochondrial and nuclear genes have been used individually and in combination to discriminate insect species [19–21]. Although multigene analysis is valuable in resolving complex taxonomic situations and is essential for phylogenetic reconstructions [22,23], it has seen little adoption for routine identifications [18]. By contrast, DNA barcoding [24] employs a 658 bp segment of a single gene, cytochrome *c* oxidase I, to discriminate animal species. Because of its ease of application and low cost, DNA barcoding has gained broad uptake [25–28]. It is now commonly used to identify specimens and to resolve sibling species complexes in insects including aphids [29–32].

The application of DNA barcoding requires bioinformatics support and a well-parameterized reference library [33]. The Barcode of Life Data System (BOLD – www.boldsystems.org) [34] meets the former need and currently includes more than six million barcode records from animals. Most of these records are from insects (5.2 million) and 49,000 records derive from aphids (accessed 3 July 2019). All barcode sequences meeting quality criteria receive a Barcode Index Number (BIN) [35]. BINs are an effective species proxy because they correspond closely with species designated through morphological study [36,37]. As a result, BINs are now routinely employed for biodiversity assessments, counting species, analyzing cryptic species complexes, and assessing species ranges [38–40]. These developments have generated a high level of interest in DNA barcoding, leading to reference barcode libraries for certain groups at continental and global scales [32,41–45].

Although the DNA barcode library for insects is incomplete, it is already valuable for identifying pest species and assessing their distributions [29,42,46–49]. However, the lack of reference sequences constrains the utility of DNA barcoding in many situations. Although barcode coverage for the aphid fauna of some countries is extensive [43,50,51] work in other nations, including Pakistan, has been limited. The current study addresses this gap by generating a barcode reference library for the pest aphids of Pakistan, and by using BINs to reveal their links to aphid assemblages in other regions.

## Materials and Methods

### Ethics Statement

No specific permissions were required for this study. The study did not involve endangered or protected species.

Aphids were sampled from 123 plant species representing 33 families at 87 sites in Pakistan (Fig. 1, S1 Table) during 2010-2013. These sites included agricultural settings, nurseries, national parks, botanical gardens, natural forests, and disturbed habitats. Based on GPS coordinates, the collection sites were rendered using SimpleMappr.net (Fig. 1). Aphids were collected by either beating foliage above a white paper sheet or by removing them from their host plant with a camel hair brush [52]. Collections were transferred into Eppendorf tubes prefilled with 95% ethanol and stored at −20°C until analysis.

**Figure 1.**
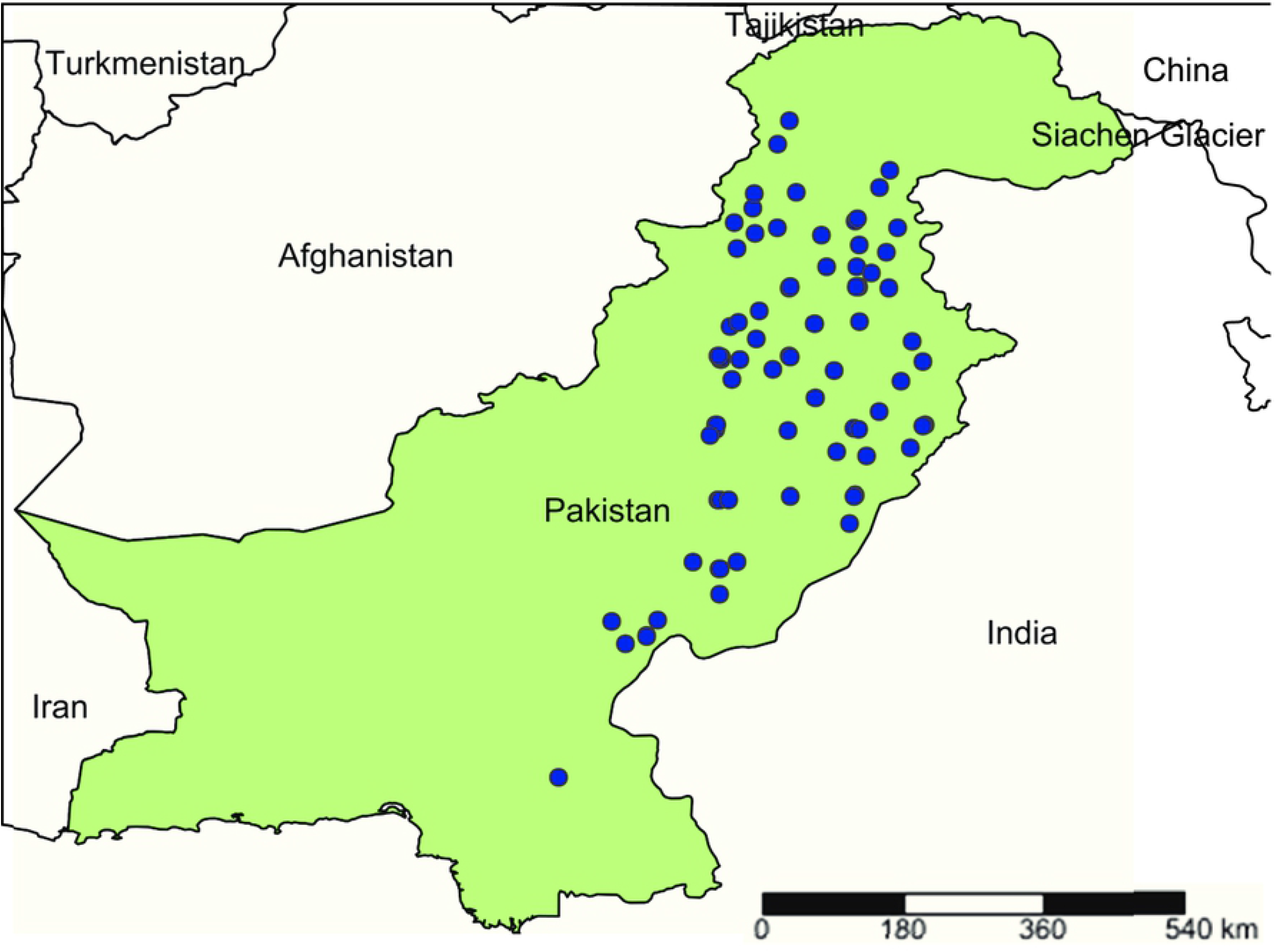
Collection sites for aphids in Pakistan. The map was generated by www.simplemappr.net using GPS coordinates.

### Identification

Aphids were identified using standard taxonomic keys [52,53]. Morphological characters were examined with a Labomed CZM6 stereomicroscope (Labo America) equipped with an ocular micrometer. Each specimen was assigned to a Linnaean species based on morphology and the identification was later validated by DNA barcode sequence matches on BOLD.

### DNA Barcoding

Front-end processing, including specimen sorting, arraying, databasing, and imaging was performed at the Insect Molecular Biology Laboratory, National Institute for Biotechnology and Genetic Engineering (NIBGE), Faisalabad. Individual specimens were placed into 96-well format in a microplate pre-filled with 30 μl of 95% ethanol in each well. Each specimen was photographed dorsally using a 12 megapixel Olympus μ-9000 camera (Olympus America Inc., USA) mounted on a stereomicroscope. Specimen metadata (collection information and taxonomy) and images were submitted to BOLD under the project MAAPH, “Barcoding Aphid Species of Pakistan”. DNA extraction, PCR amplification, and sequencing were carried out at Centre for Biodiversity Genomics at Guelph. DNA extraction followed Ivanova et al. [54] with voucher recovery protocol [55]. PCR amplification of the COI-5′ (barcode region) [24] was performed using primer pair C_LepFolF (forward) and C_LepFolR (reverse) (http://ccdb.ca/site/wp-content/uploads/2016/09/CCDB_PrimerSets.pdf) in 12.5 μL reactions that included standard PCR ingredients [56] and 2 μL of DNA template. The thermocycling regime was: 94°C (1 min), 5 cycles at 94°C (40 s), 45°C (40 s), 72°C (1 min); 35 cycles at 94°C (40 s), 51°C (40 s), 72°C (1 min); and a final extension at 72°C (5 min). PCR success was verified by analyzing the amplicons on 2% agarose E-gel^®^ 96 system (Invitrogen Inc.). Specimens which failed to amplify in the first round of PCR were re-run with primers LepF2_t1 (TGTAAAACGACGGCCAGTAATCATAARGATATYGG) [57] and LepR1 using the same PCR conditions. PCR products were sequenced bidirectionally on an Applied Biosystems 3730XL DNA Analyzer using the BigDye Terminator Cycle Sequencing Kit (v3.1) (Applied Biosystems). Sequences were edited using CodonCode Aligner (CodonCode Corporation, USA), and translated on MEGA v6 [58] to confirm they were free of stop codons, and submitted to BOLD. The specimen metadata and sequences generated in this study are available on BOLD in the dataset DS-MAAPH.

### Data Analysis

All barcode sequences were compared with those on BOLD and GenBank to ascertain sequence similarities. Sequence matches on BOLD were obtained using “Identification Engine” while nBLAST (http://www.ncbi.nlm.nih.gov/blast/) was used on GenBank. All sequences meeting quality standards (>500 bp, <1% ambiguous bases, no stop codon or contamination flag) were assigned to a BIN [35]. BIN discordance and BIN overlap reports were generated using analytical tools on BOLD. As a test of the reliability of species discrimination, the presence or absence of a ‘barcode gap’ [59] among species was determined on BOLD. A species was considered distinct when its maximum intraspecific distance was less than the distance to its nearest neighbor (NN).

Nucleotide alignments and neighbor-joining (NJ) analysis [60] were conducted in MEGA6. The NJ analysis employed the Kimura-2-Parameter (K2P) [61] distance model, with pairwise deletion of missing sites, and 1000 bootstrap replicates for the nodal support. Bayesian inference was performed in MrBayes v3.2.0 [62] employing the Markov Chain Monte Carlo (MCMC) technique. This analysis was based on representative sequences from 67 aphid haplotypes in the dataset extracted using DNaSP v5.10 [63] with *Diaphorina citri* (Hemiptera: Psyllidae) as outgroup. The data were partitioned in two ways; i) a single partition with parameters estimated across all codon positions, ii) a codon-partition in which each codon position was allowed different parameter estimates. The evolution of sequences was modelled by the GTR+Γ model independently for the two partitions using the “unlink” command in MrBayes, and the best model was selected using FindModel (www.hiv.lanl.gov/cgi-bin/findmodel/findmodel.cgi). The analyses were run for 10 million generations using four chains with sampling every 1000 generations. Bayesian posterior probabilities were calculated from the sample points once the MCMC algorithm converged. Convergence was defined as the point where the standard deviation of split frequencies was less than 0.002 and the PSRF (potential scale reduction factor) approached 1, and both runs converged to a stationary distribution after the burn-in (by default, the first 25% of samples were discarded). Each run produced 100001 samples of which 75001 samples were included. The trees generated through this process were visualized using FigTree v1.4.0.

BOLD was searched for barcode records for the 42 Linnaean species encountered in this study. The resultant records were examined for BIN assignment [35] and used in a geo-distance correlation analysis to examine the relationship between geographic and genetic distance in each species. Two methods were employed in the latter analysis; the Mantel Test [64] was used to examine the relationship between geographic (km) and genetic (K2P) distance matrices, while the second approach compared the spread of the minimum spanning tree of collection sites and maximum intra-specific divergence [65]. The relationship between geographic and intraspecific distances was analyzed for each species with at least one individual from three or more sites. BINs recovered from Pakistan were also used in BIN-overlap analysis on BOLD to ascertain the incidence of unique BINs in a region, and to estimate overlap in BIN composition.

## Results

Morphological analysis facilitated by the barcode data allowed 774 of the 809 specimens to be identified to 42 species, each an important crop pest (S1 Table). Another 32 specimens could be placed to a genus while the other three could only be assigned to a subfamily (Aphidinae). Overall, the 809 specimens included representatives of 30 genera, five subfamilies (Aphidinae, Calaphidinae, Chaitophorinae, Eriosomatinae, Lachninae) of the Aphididae (S2 Table). Members of the Aphidinae were dominant (n=780) as the other four subfamilies were represented by just 29 specimens with Chaitophorinae and Lachninae each contributing one specimen (S2 Table). Among the genera, *Aphis* was most common (n=306), and it was represented by eight identified and three undetermined species. *Myzus* was the second most frequent genus (n=170), but it was only represented by one species, *Myzus persicae. Rhopalosiphum*, the third most abundant (83) genus, was represented by three major pest species (*R. maidis, R. padi, R. rufiabdominale*). Two species (*Aphis astragalina, Periphyllus lyropictus*) represented first records for Pakistan while two others (*Lipaphis pseudobrassicae, Sarucallis kahawaluokalani*) were known, but were recorded as *Lipaphis erysimi* and *Tinocallis kahawaluokalani*, names now relegated to synonymy.

The 809 barcode sequences provided two or more records for 36 of the 42 species and single records for the rest (Table 1, S1). Maximum K2P divergence values within species ranged from 0 – 3.6% (mean=0.1%), while the within genus values were 0.8 – 10.3% (mean=7.4%), and within family (Aphididae) 3.7 – 17.3% (mean=9.6%) (Table 1). Barcode gap analysis examined the ability of barcodes to discriminate the 42 named species. With the exception of one species (*Aphis gossypii*), where the maximum intraspecific distance (3.6%) overlapped with *A. affinis*, the maximum intraspecific distance for each species was less than its NN distance (Fig. 2A + Fig. 2B). This pattern did not change with increased sample size (Fig. 2C).

**Figure 2.**
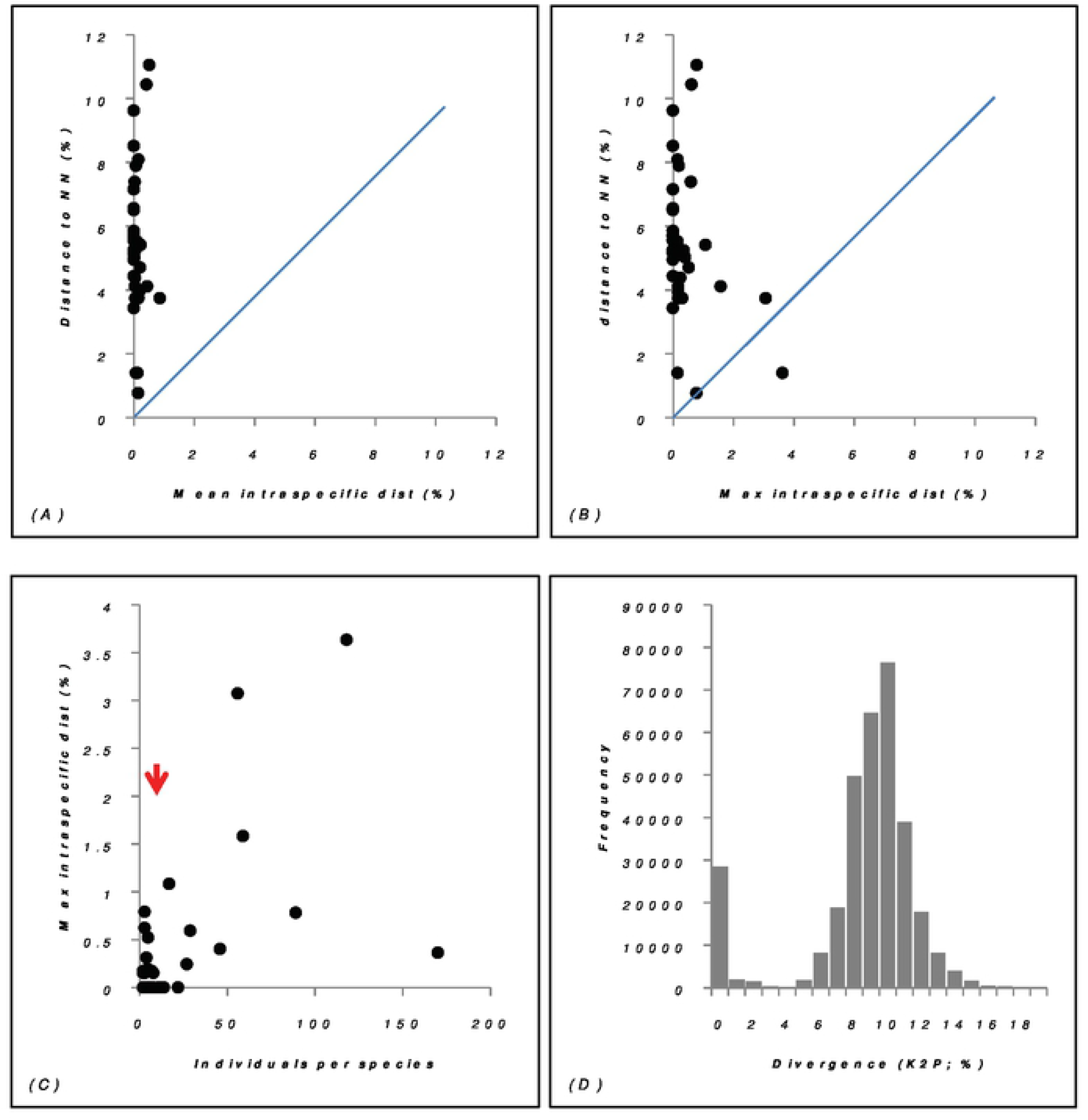
Barcode gap analysis for species of aphids with three or more specimens collected in Pakistan. NN = nearest neighbor.

**Table 1.** Sequence divergence (K2P) in the COI barcode region for aphid species from Pakistan with more than three specimens, genera with three or more species, and families with three or more genera. This analysis only considers specimens that were assigned to a Linnaean species.

| Distance class | n | Taxa | Comparisons | Min (%) | Mean (%) | Max (%) |
| --- | --- | --- | --- | --- | --- | --- |
| Within Species | 764 | 36 | 30756 | 0 | 0.1 | 3.6 |
| Within Genus | 434 | 6 | 32305 | 0.8 | 7.4 | 10.3 |
| Within Family | 770 | 1 | 233004 | 3.7 | 9.6 | 17.3 |

Nearly all sequences (801/809) qualified for a BIN assignment, and they were placed in 52 BINs. The 774 specimens of the 42 Linnaean species were assigned to 41 BINs; 38 showed BIN concordance (species members = BIN members), one species (*Rhopalosiphum padi*) was split (AAA9899, ACF2924), and two species (*Aphis affinis, A. gossypii*) were merged (AAA3070), while another (*Aphis astragalina*) lacked a BIN assignment due to its low quality sequence (410 bp, 9 Ns). The 32 specimens lacking a species assignment were placed in 9 BINs – three for *Aphis* and one for each of the other six genera (*Acyrthosiphon, Capitophorus, Forda, Hyalopterus, Macrosiphoniella, Schizaphis*). The three specimens only identified to a subfamily were assigned to two BINs. NJ analysis (Fig. 3) and Bayesian inference (BI) (Fig. 4) supported the monophyly of each of the 52 BINs. The NJ and BI also discriminated the species or genera that either lacked (*Aphis astragalina*) or shared BINs (*Aphis gossypii*, *A. affinis*), as they formed distinct branches on the NJ and BI trees (Fig. 3, Fig. 4).

**Figure 3.**
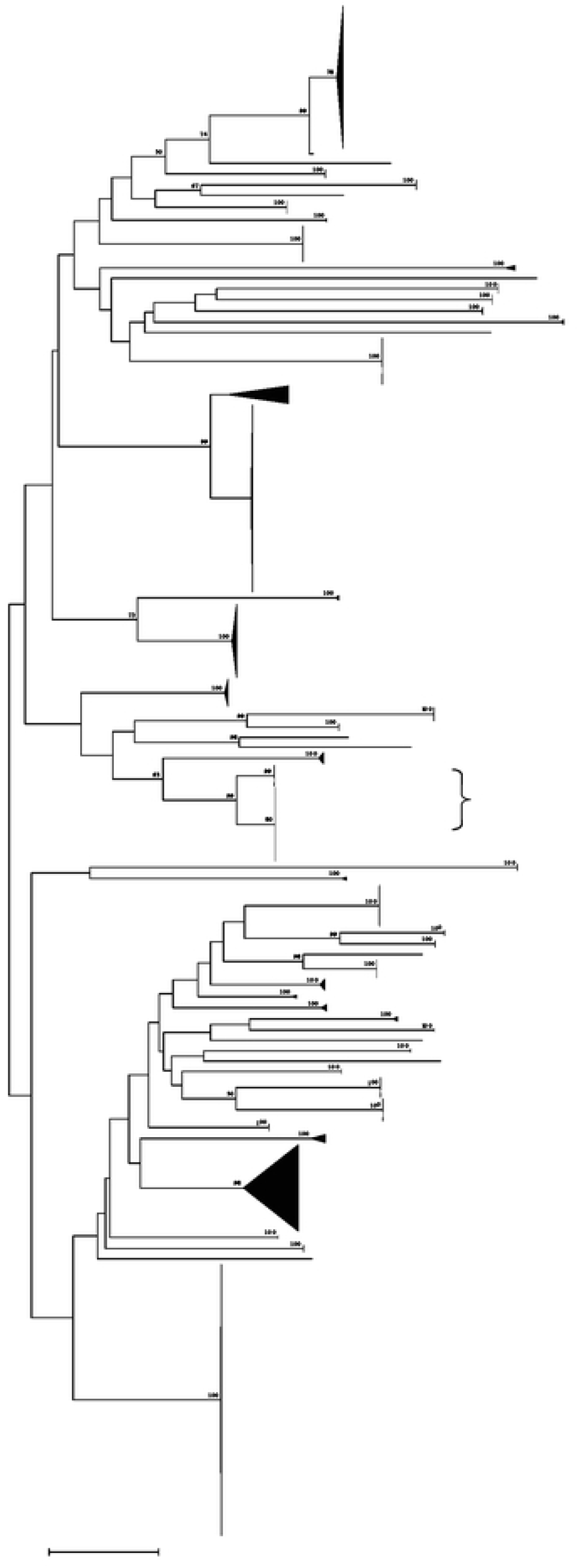
NJ analysis of COI-5′ sequences from species/ BINs of aphids from Pakistan. Bootstrap values (%) (1,000 replicates) are shown above the branches (values <50% are not shown) while the scale bar shows K2P distances. The node for each species/BIN with multiple specimens was collapsed to a vertical line or triangle, with the horizontal depth indicating the level of intraspecific divergence. BIN numbers are shown for species with only family- or genus-level identification or those split into two BINs.

**Figure 4.**
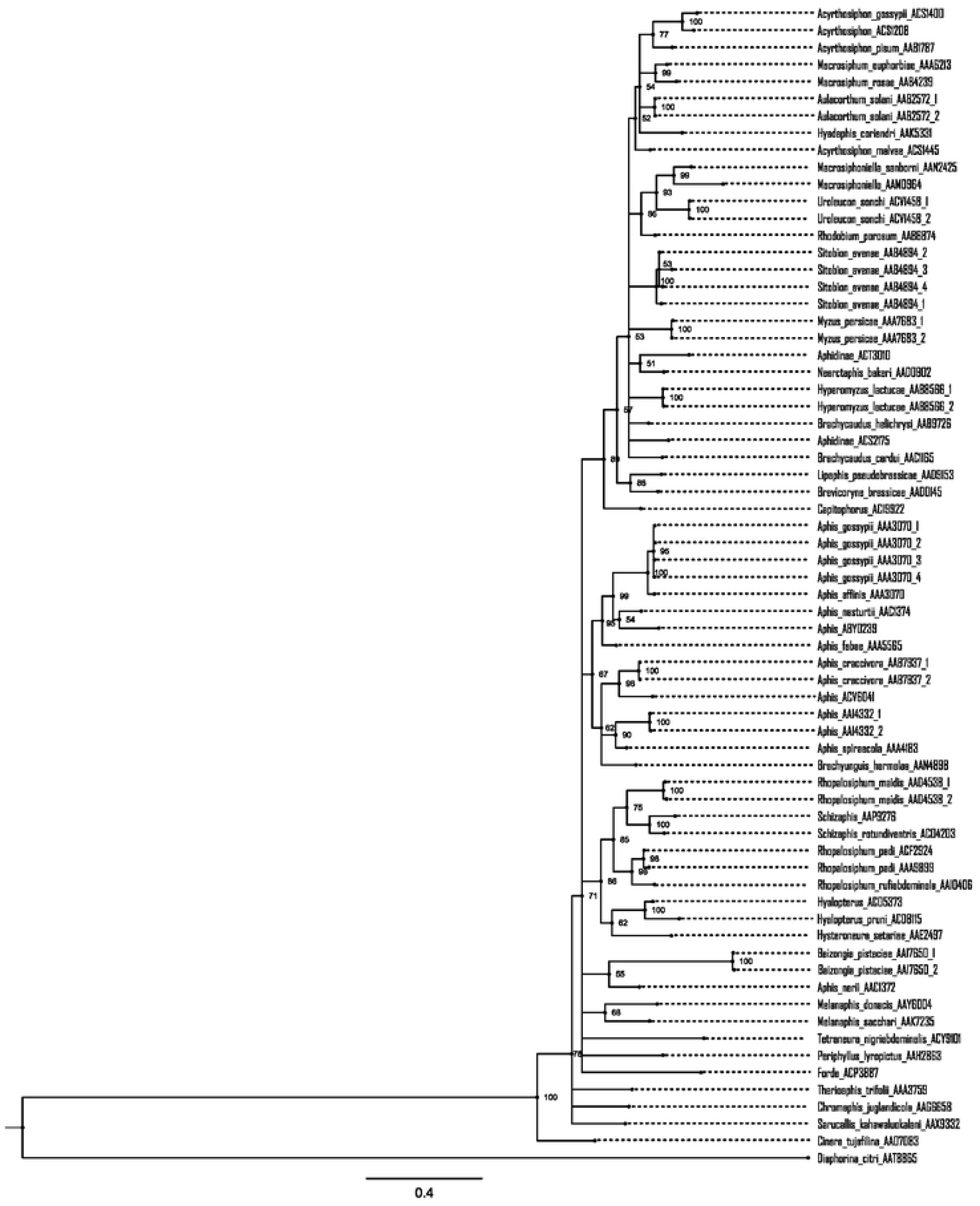
Phylogenetic analysis of aphid species/BINs from Pakistan based on COI-5’ sequences. The tree was estimated using Bayesian inference. Posterior probabilities are indicated at nodes. The analysis was based on representative sequences from 67 aphid haplotypes in the dataset that were extracted using DnaSP v5.10 (Librado and Rozas 2009). Taxa are followed by the BINs and haplotype numbers. *Diaphorina citri* (BOLD:AAT8865) was employed as outgroup.

Geo-distance correlation analysis for 37 species was conducted by including 5,067 sequences from conspecific individuals on BOLD. This analysis showed that intraspecific divergence in 49% of the species was not affected by expanding analysis to consider its entire range (Mantel test, P>0.01) (Table 2). The other 51%, that were affected by geographic range, included six species with BIN splits and eight with intraspecific divergence higher than >2%. The distributional patterns of aphids detected in Pakistan were further analyzed by examining BIN overlap between Pakistan and other countries, a comparison that involved 9,905 barcode records assigned to the 52 BINs. This analysis showed that 27 of the 52 BINs were recorded from four or more continents while eight were unique to Pakistan (Table 3). Except for *Acyrthosiphon malvae* and *Uroleucon sonchi*, all named species (40) analyzed in this study already had barcode records from multiple countries and continents (Table 3).

**Table 2:** Geographic (km) and genetic (K2P) distance correlation analysis for 42 aphid species from Pakistan combined with conspecifics from 69 other countries.

| Species | Record Count | BINs | Linear Regression R<sup>2</sup> | Gen Dist Max | Geo Dist Max | Mantel R<sup>2</sup> | Mantel P value |
| --- | --- | --- | --- | --- | --- | --- | --- |
| *Acyrtosiphon malvae* | 54 | 2 | 0.13 | 4.8 | 18675 | 0.13 | 0.01 |
| *Acyrtosiphon pisum* | 205 | 1 | 0.01 | 1.7 | 19312 | 0.01 | 0.05 |
| *Aphis affinis* | 8 | 1 | 0.18 | 0.2 | 356 | 0.18 | 0.04 |
| *Aphis astragalina* | 37 | 1 | 0.71 | 0.8 | 10789 | 0.71 | 0.01 |
| *Aphis craccivora* | 420 | 1 | 0.09 | 1.4 | 19417 | 0.00 | 0.01 |
| *Aphis fabae* | 426 | 1 | 0.01 | 2.5 | 19456 | 0.01 | 0.01 |
| *Aphis gossypii* | 362 | 1 | 0.00 | 3.9 | 19369 | 0.00 | 0.8 |
| *Aphis nasturtii* | 38 | 1 | 0.48 | 1.6 | 11656 | 0.48 | 0.01 |
| *Aphis nerii* | 99 | 1 | 0.02 | 1.2 | 19110 | 0.02 | 0.01 |
| *Aphis spiraeicola* | 277 | 2 | 0.00 | 3.1 | 19355 | 0.00 | 0.2 |
| *Aulacorthum solani* | 118 | 1 | 0.13 | 2.0 | 19291 | 0.13 | 0.01 |
| *Baizongia pistaciae* | 5 | 2 | 0.48 | 4.4 | 5967 | 0.48 | 0.23 |
| *Brachycaudus cardui* | 54 | 1 | 0.17 | 0.9 | 11929 | 0.17 | 0.01 |
| *Brachycaudus helichrysi* | 108 | 4 | 0.01 | 3.1 | 19426 | 0.01 | 0.01 |
| *Brevicoryne brassicae* | 166 | 2 | 0.02 | 6.3 | 19178 | 0.02 | 0.04 |
| *Chromaphis juglandicola* | 11 | 1 | 0.52 | 0.8 | 10908 | 0.52 | 0.01 |
| *Hyadaphis coriandri* | 13 | 1 | 0.40 | 1.4 | 685 | 0.40 | 0.01 |
| *Hyalopterus pruni* | 151 | 3 | 0.06 | 6.6 | 16867 | 0.06 | 0.01 |
| *Hyperomyzus lactucae* | 87 | 1 | 0.05 | 0.3 | 19474 | 0.05 | 0.02 |
| *Hysteroneura setariae* | 53 | 1 | 0.04 | 1.9 | 15843 | 0.04 | 0.01 |
| *Lipaphis pseudobrassicae* | 91 | 2 | 0.12 | 3.6 | 18212 | 0.13 | 0.01 |
| *Macrosiphoniella sanborni* | 5 | 1 | 0.01 | 0.2 | 15935 | 0.01 | 0.43 |
| *Macrosiphum euphorbiae* | 198 | 1 | 0.14 | 1.6 | 19482 | 0.14 | 0.01 |
| *Macrosiphum rosae* | 80 | 1 | 0.00 | 1.2 | 19379 | 0.00 | 0.14 |
| *Melanaphis donacis* | 7 | 1 | 0.16 | 0.2 | 6092 | 0.16 | 0.35 |
| *Melanaphis sacchari* | 225 | 1 | 0.02 | 1.9 | 18400 | 0.02 | 0.01 |
| *Myzus persicae* | 322 | 1 | 0.00 | 2.2 | 19234 | 0.00 | 0.68 |
| *Nearctaphis bakeri* | 70 | 2 | 0.04 | 6.8 | 11961 | 0.04 | 0.09 |
| *Periphyllus lyropictus* | 33 | 1 | 0.00 | 0.2 | 11002 | 0.00 | 0.92 |
| *Rhodobium porosum* | 7 | 1 | 0.08 | 0.2 | 12350 | 0.08 | 0.1 |
| *Rhopalosiphum maidis* | 63 | 1 | 0.00 | 2.0 | 18405 | 0.00 | 0.40 |
| *Rhopalosiphum padi* | 1189 | 2 | 0.30 | 5.0 | 19841 | 0.29 | 0.01 |
| *Rhopalosiphum rufiabdominale* | 18 | 1 | 0.03 | 0.2 | 15787 | 0.03 | 0.99 |
| *Sitobion avenae* | 314 | 1 | 0.02 | 4.6 | 18405 | 0.00 | 0.01 |
| *Tetraneura nigriabdominalis* | 41 | 2 | 0.16 | 8.6 | 16534 | 0.16 | 0.01 |
| *Therioaphis trifolii* | 470 | 2 | 0.00 | 13.0 | 18823 | 0.00 | 0.33 |
| *Uroleucon sonchi* | 45 | 2 | 0.11 | 2.3 | 19003 | 0.11 | 0.03 |
| *Acyrtosiphon gossypii* | 5 | 1 | N/A | N/A | N/A | N/A | N/A |
| *Brachyunguis harmalae* | 8 | 1 | N/A | N/A | N/A | N/A | N/A |
| *Cinara tujafilina* | 8 | 1 | N/A | N/A | N/A | N/A | N/A |
| *Sarucallis kahawaluokalani* | 10 | 1 | N/A | N/A | N/A | N/A | N/A |
| *Schizaphis rotundiventris* | 5 | 1 | N/A | N/A | N/A | N/A | N/A |

N/A: Data for the correlation analysis was missing.

**Table 3:**
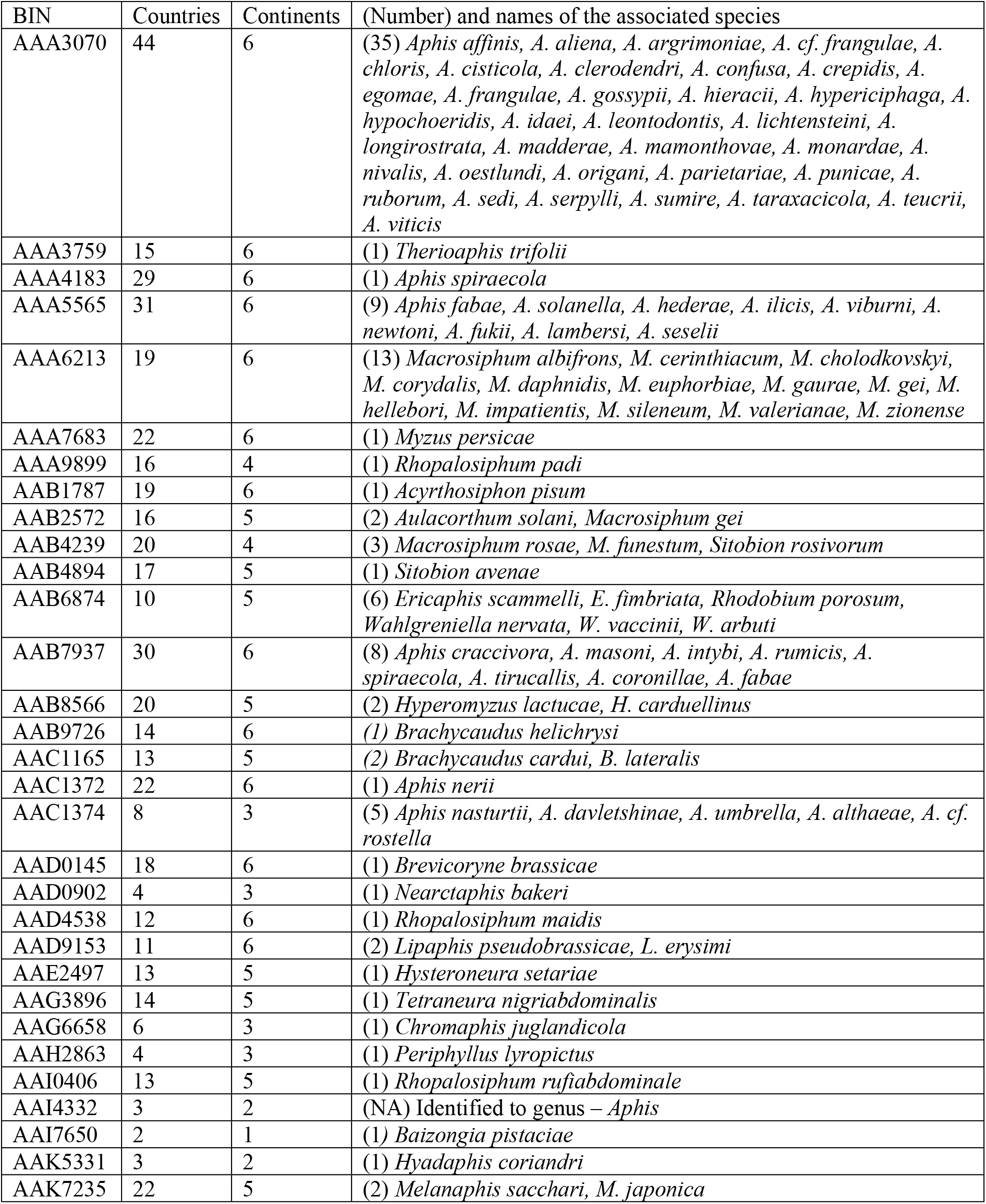
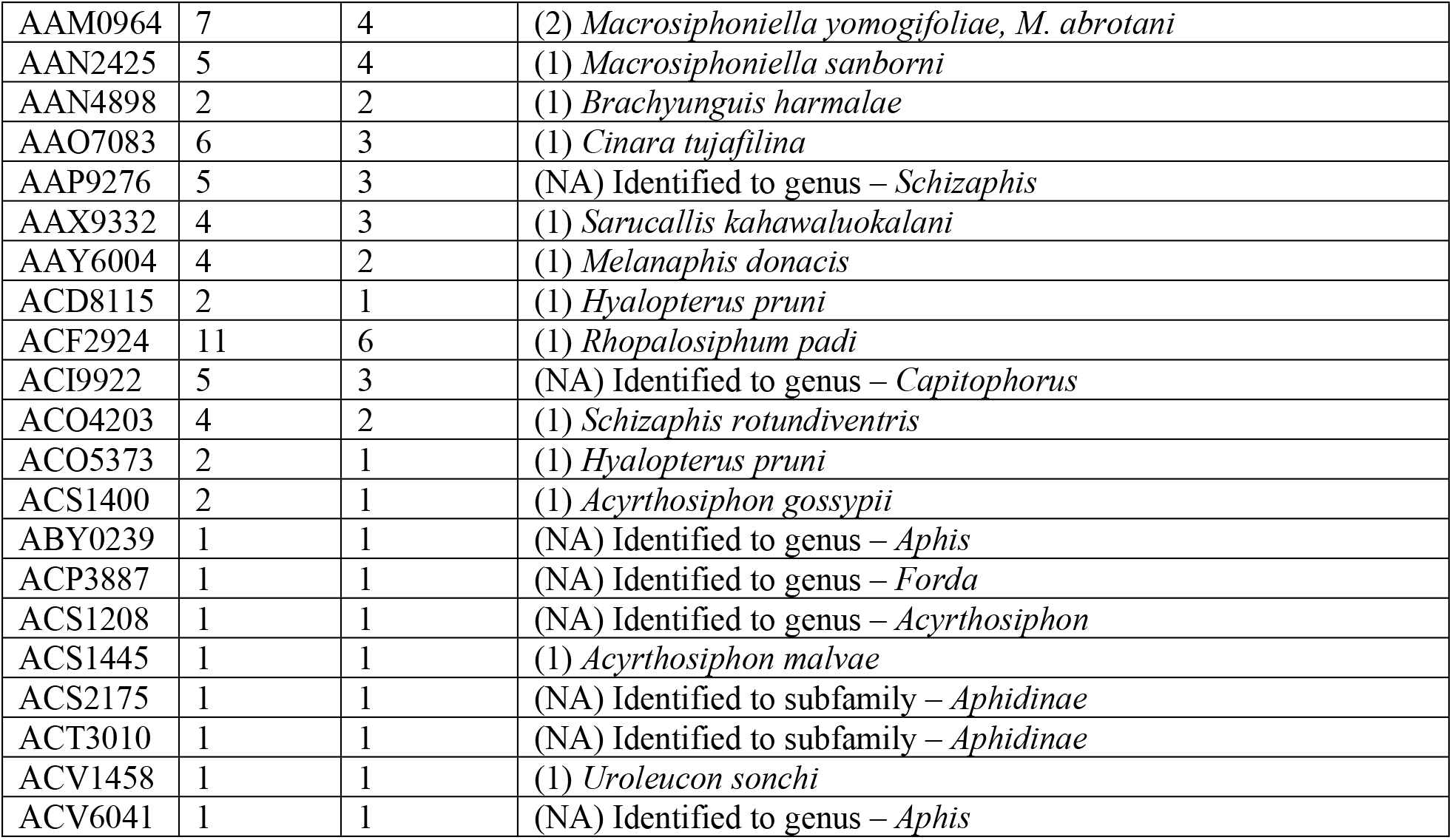
Occurrence of 52 pest aphid BINs across six continents and their association with Linnaean species on the Barcode of Life Data System (BOLD).

## Discussion

Prior morphological surveys on the aphids of Pakistan have reported the presence of nearly 300 species [66–68]. Most of this work focused on specific geographic regions [69] or on species attacking crops [70,71]. The current study surveyed aphids across much of Pakistan from a wider range of host plants, but primarily aimed to develop a barcode reference library for the fauna. Prior studies have begun to construct barcode reference libraries for some pest insect groups, such as aphids in Canada [29], leafminers in USA [72], fruit flies in Africa [42], food pests in Korea [46], thrips in Pakistan [28], looper moths in British Columbia [73], and mealybugs in China [49]. These libraries have stimulated the use of DNA barcoding in biosecurity and plant protection programs [74], but this use have revealed the need for expanded parameterization of the libraries to improve their utility in diagnosing newly encountered species. Barcode libraries for two major pest insect groups in Pakistan, thrips and whiteflies, are well advanced [28,75], but other groups have seen little attention in this country.

Most aphids analyzed in this study could be assigned to a species, but 35 specimens could only be resolved to a genus or subfamily. In part, this difficulty reflected the fact that many important pest aphids are cryptic species complexes whose members are almost impossible to discriminate morphologically [39]. For example, *Aphis gossypii* is a particularly challenging species complex [5,13]; it includes at least 20 morphologically indistinguishable species [76] likely explaining its wide range of primary and secondary host plants [77]. In the present study, DNA barcoding separated all eight species of the genus *Aphis* that were encountered. Although K2P distances between two species pairs; i) *A. affinis* and *A. gossypii* (1.4%), ii) *A. astragalina* and *A. craccivora* (0.8%) were low, both NJ analysis and Bayesian inference supported the monophyly of each species. The COI divergences in this study are similar to those reported in prior investigations [29,78,79] which reported low sequence divergence between sibling species such as *A. gossypii* and *A. affinis* [29].

Prior studies have shown strong correspondence between BINs and known species [36], especially when reference specimens are identified by experts [80]. The same pattern was apparent in this study as 38 of 41 Linnaean species were assigned to a single BIN. There were only two exceptions; *R. padi* was assigned to two BINs and *A. gossypii* – *A. affinis* were assigned to the same BIN. By comparison, when barcode sequences from conspecific specimens from other countries were considered, 12 of the 4 species showed a BIN split, an outcome which likely indicates incorrectly identified specimens [36]. Interestingly, the BIN (AAA3070) shared by specimens of *A. gossypii* and *A. affinis* from Pakistan included 31 additional species names when all records for it on BOLD were considered. Misidentifications and overlooked cryptic species may often cause conflicts between BIN and species morphology [81], but this can only be resolved by detailed taxonomic studies [82].

Geo-distance correlations showed that the genetic divergence in almost half of the aphid species increased with geographic distance while the others were unimpacted. Interestingly, the inclusion of conspecific sequences from other regions also increased the incidence of BIN splits. Since these analyses included all the conspecific sequences on BOLD, this outcome may reflect taxonomic errors [83]. Although spatial variation in conspecific sequences sometimes leads to increased intraspecific divergence values [84], it is usually too low to reduce the capacity of DNA barcodes to deliver reliable species identifications [44,85].

BINs are valuable in evaluating the geographic range of aphid species because they circumvent taxonomic uncertainties. BINs are gaining increased use to estimate species numbers [38] and to understand their distributions [49]. This analysis revealed that 27 of the 44 BINs with prior records on BOLD occurred on four or more continents, highlighting the broad ranges of many pest aphids. For example, BINs for *Aphis fabae* (black bean aphid), *A. nerii* (oleander aphid), *A. craccivora* (groundnut aphid), *Acyrthosiphon pisum* (pea aphid), *Brachycaudus helichrysi* (plum aphid), *Brevicoryne brassicae* (cabbage aphid), *L. pseudobrassicae* (turnip aphid), *R. padi* (oat aphid), *R. maidis* (corn aphid), *Macrosiphum euphorbiae* (potato aphid), *M. persicae* (peach aphid), and *Therioaphis trifolii* (alfalfa aphid) were all recorded from six continents. Interestingly, BINs associated with some of these species were also linked with other species on BOLD. For instance, AAA3070 was linked to 33 other species of *Aphis* while AAA6213 was associated with 13 species of *Macrosiphum*, and AAA5565 with nine species of *Aphis*. Although some of these cases may involve BIN sharing by different species [29], most cases likely reflect misidentifications.

The level of BIN overlap between the aphid fauna of Pakistan is much higher (85%) than levels for moths (44%) [86] and spiders (24%) [87]. This difference, may, be due, in part, to the fact that the winged alates of aphids can disperse long distances are they use widely crop plants as their hosts [88]. Consequently, the number of aphid species known from Europe has increased by 20% in the last 30 years [89] reflecting their transport on produce fruits [49], coupled with shifting environmental regimes. Reports suggest that with every 1°C increase, some 15 additional aphid species were recorded in Europe [90]. In North America, about 18% of all aphid species are introduced, and nearly half are plant pests [91]. Rapid developments in DNA sequencing are enabling the documentation of pest species and their distribution across the globe, but conflicts between taxonomic assignments and sequences have limited the full utility of these data. Given this difficulty, the BIN system provides an alternative path to document and track the pest species on a planetary scale.

## Acknowledgments

We are very grateful to C Favret, University of Montreal, Quebec, Canada, for his help in aphid identification. We thank staff at the CCDB for aiding the sequence analysis which was made possible by a grant from the Government of Canada through Genome Canada and Ontario Genomics in support of the International Barcode of Life (iBOL) project.

## Supporting Information Legends

S1 Table. Plant-host family range for aphid species/BINs collected in Pakistan.

S2 Table. Identification and BIN assignment of 809 specimens of Aphididae collected from 123 plant hosts in Pakistan.

